# Thermal Adaptation of Cytosolic Malate Dehydrogenase Revealed by Deep Learning and Coevolutionary Analysis

**DOI:** 10.1101/2024.10.08.617074

**Authors:** D Shukla, J Martin, F Morcos, DA Potoyan

## Abstract

Protein evolution has produced enzymes that maintain stability and function across various thermal environments. While sequence variation, structural dynamics, and intermolecular interactions are known to influence an enzyme’s thermal adaptation, how these factors collectively govern stability and function across diverse temperatures remains unresolved. Cytosolic malate dehydrogenase (cMDH), a citric acid cycle enzyme, is an ideal model for studying these mechanisms due to its temperature-sensitive flexibility and broad presence in species from diverse thermal environments. In this study, we employ techniques inspired by deep learning and statistical mechanics to uncover how sequence variation and structural dynamics shape patterns of cMDH’s thermal adaptation. By integrating coevolutionary models with variational autoencoders (VAE), we generate a latent generative landscape (LGL) of cMDH sequence space, enabling us to explore evolutionary pathways and predict fitness using direct coupling analysis (DCA). Structural predictions via AlphaFold and molecular dynamics simulations further illuminate how variations in hydrophobic interactions and conformational flexibility contribute to the thermal stability of warm- and cold-adapted cMDH orthologs. The integrative computational framework employed in this study provides powerful insights into protein adaptation at both sequence and structural levels, offering new perspectives on the evolution of thermal stability and creating avenues for the rational design of proteins with optimized thermal properties for biotechnological applications.

## Introduction

Enzymes from different organisms have evolved to maintain an optimal balance of stability and functional activity under various thermal environments. While variations in sequence, structural dynamics, and intermolecular interactions influence an enzyme’s thermal adaptation, the molecular rules that collectively govern stability and function across diverse temperatures remain unclear. Cytosolic malate dehydrogenase (cMDH), a well-characterized enzyme in the citric acid cycle, is an exemplary model system for investigating these mechanisms due to its temperature-sensitive conformational flexibility and widespread occurrence in species adapted to various thermal niches [1–4]. The homologs of cMDH display an impressive range of thermal adaption ranging from approximately -20°C to over 120°C.[5,6] The wealth of information collected for cMDH enables detailed studies of specific sites within the sequence critical for ligand binding, catalysis, and subunit interactions. The conformational changes necessary for cMDH catalytic function are well understood, aiding the structure-function analyses [2]. A recent study[7] has identified key mobile regions (MRs) that undergo significant conformational changes, enabling ligand binding and catalytic activity. Dynamics of Cytosolic MDH from orthologs from warm-adapted and cold-adapted species display distinct thermal responses[5,8]. For instance, cytosolic malate dehydrogenase (cMDH) from warm-adapted *Mytilus galloprovincialis* maintains activity and optimal substrate binding at higher temperatures than their cold-adapted counterparts from *Mytilus trossulus*. This is mainly due to amino acid substitutions that enhance structural stability[5,9]

Numerous investigations have focused on comparing psychrophilic, thermophilic, and mesophilic proteins to pinpoint the molecular determinants of protein thermal stability[10–12]. Changes in non-covalent intramolecular interactions, including electrostatic interactions, hydrogen bonds, and hydrophobic contacts, have been found to correlate with varying thermal stabilities observed among homologous proteins [13,14]. Evolutionary changes, however, are known to be constrained by the fitness of an organism, which exerts selective pressures on the sequence, structure, and dynamics of proteins [15–17]. These constraints impose statistical signatures in the collection of evolutionarily related sequences, allowing structural and functional inferences from homologous sequence alignments, which is exploited in techniques collectively known as direct coupling analysis (DCA) [18–20]. The inference of co-evolutionary pairs in DCA has shown to be crucial for the success of next-generation AI-based structure prediction techniques of AlphaFold and RoesTTA, which can reconstruct 3D folds of single proteins and [22–24] protein complexes [21–24] from multiple sequence alignments. Co-evolutionary models have also proven to be key in inferring molecular specificity and the effects of protein mutations, thereby guiding the design of novel functional proteins such as transporters, fluorescent proteins, and enzymes[18,20,25]. Recent focus has shifted towards using state-of-the-art AI approaches in extending the predictive range of co-evolutionary models towards functional relationships in protein families[26,27]. For instance, DeepPPI has been used to study protein interactions, demonstrating its effectiveness in identifying interaction networks [28]. Restricted Boltzmann machines (RBM) have been applied to detect motifs associated with functional roles in proteins, highlighting their utility in motif discovery [29]. Architectures such as Variational autoencoders (VAE) and Transformers have shown especially high promise in generating novel protein sequences, underlining their potential in protein design [30]. VAEs have been utilized for phylogenetic clustering and predicting the effects of protein mutations[31].

In this work, we combine the predictive power of coevolutionary models with the classification and generative power of the VAE, which allows the generation and modification of cMDH sequences in a manner that requires no labeling information. Through this combination of statistical models, we create a latent generative landscape (LGL), as shown first in the work by [32], where accessible VAE sequence space is assessed using the inferred fitness from DCA. By exploring a large amount of sequence space, we have uncovered a method to traverse the diversity of sequence space of malate dehydrogenase (MDH) that is more flexible than other architectures, such as transformers and generative adversarial networks (GANs), due to higher diversity of encoded possible sequences, no required labeling, and easily accessed latent representations [32]. Using AlphaFold and MD simulations guided by LGL, we gain insights into how evolution tunes stability-dynamics balance, enabling functional diversification and potentially guiding generative enzyme design (Fig 1).

**Fig 1:**
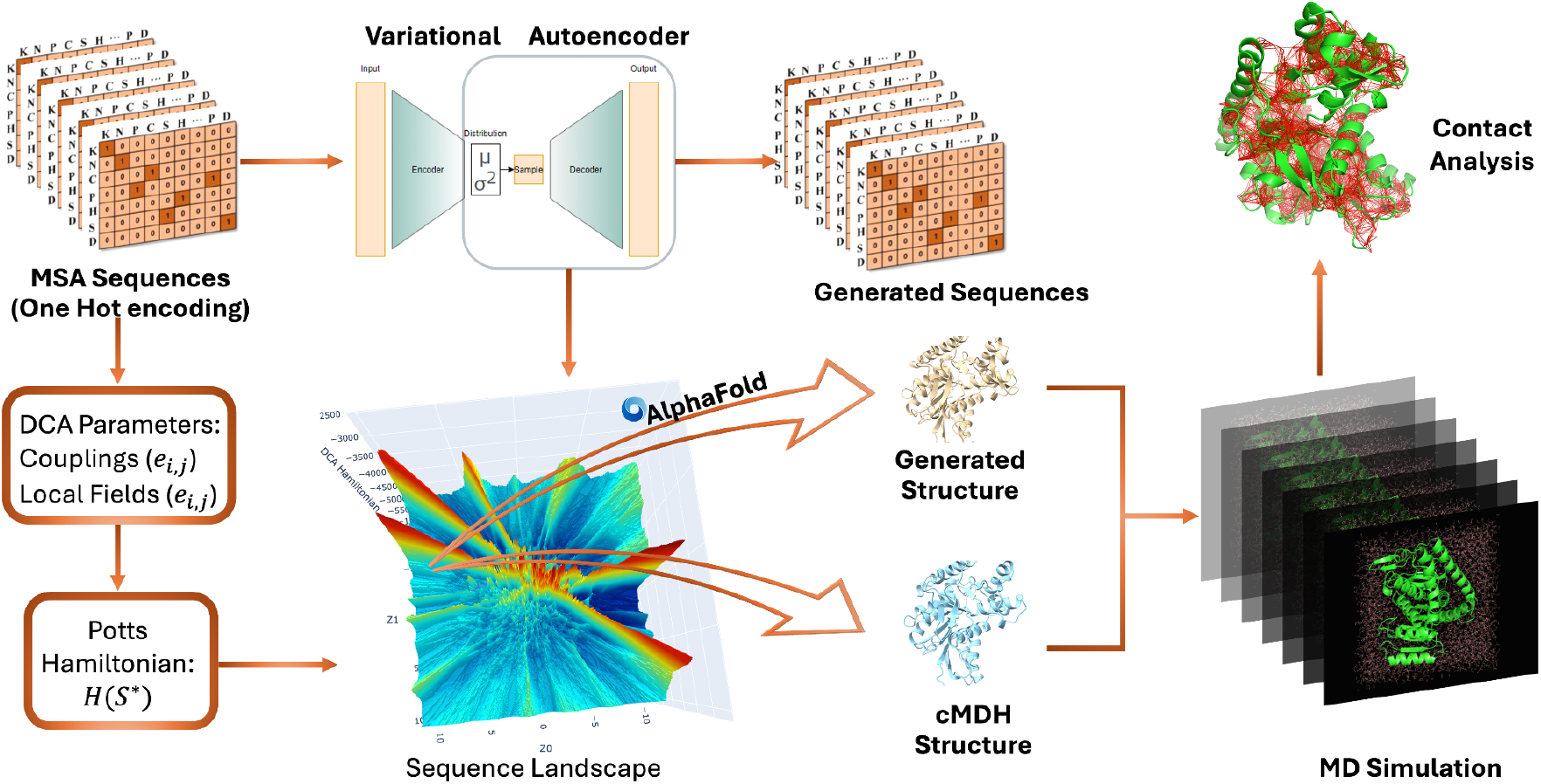
Workflow for cMDH Landscape Generation and Structural Analysis: The workflow begins with multiple sequence alignments (MSA) encoded using one-hot encoding, generating a comprehensive evolutionary landscape. These sequences are then processed through a Variational Autoencoder (VAE) to produce a two-dimensional representation of the sequence space. Fitness is assessed using a Potts Hamiltonian, with parameters derived from Direct Coupling Analysis (DCA) to evaluate coevolutionary relationships between amino acids. Predicted sequences are input into AlphaFold for three-dimensional structure prediction, including the cytosolic variant of malate dehydrogenase (cMDH). Molecular dynamics simulations are subsequently used to explore the dynamic properties of the structures, with contact analysis revealing variations in hydrophobic contact networks.

## Methods

### Generation of Multiple Sequence Alignments

For the analysis of cMDH, multiple sequence alignments (MSAs) were initially procured using the HMMSearch tool against the UniProt database, employing the GREMLIN tool developed by Baker Lab([33]; [34]). The MSAs were further refined by excluding sequences that contained more than 10% gaps, ensuring the retention of high-quality sequence data([35]). After this filtration process, a total of 2,100 sequences remained, which were compiled into a FASTA file. This curated dataset was subsequently utilized to train a Variational Autoencoder (VAE), facilitating the exploration of sequence variability and potential structural predictions([31]).

### VAE model architecture

The variational autoencoder (VAE) is designed to generate data samples *x* ϵ *X* utilizing a latent variable model defined on parameters θ with a prior p_θ_(z) on latent variables *z*. The marginal likelihood is represented as:

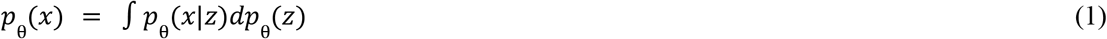

However, the parameters θ and the latent variables *z* are unknown, and Equation (1) lacks a tractable algorithmic solution for their determination. The proposed solution, as outlined in reference [36], is to approximate the posterior distribution *p*_θ_(*Z*|*x*) with another model defined on parameters Φ :

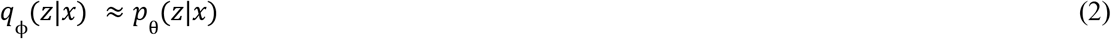

The model defined on Φ is termed the encoder, while the model defined on θ is termed the decoder. Consequently, the marginal likelihood of generating a sample *x* through the decoder can be expressed as:

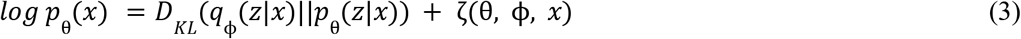

Here, *D*_*KL*_ represents the Kullback-Leibler divergence, quantifying the fit between the decoder’s posterior distribution on *z* and the encoder’s posterior, while the term ζrepresents the lower bound of the model’s fit to the marginal distribution over *z*. Equation (3) can be further simplified into the evidence lower bound function (ELBO):

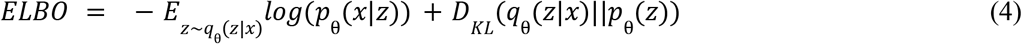

This serves as the objective function minimized during training. The reconstruction error, measuring the match between encoded and generated data, constitutes the first term, while the second term evaluates the fit of the latent distribution being encoded and an assumed prior distribution. We employ the reparameterization procedure [36] to encode the posterior to ensure differentiability. The encoder model represents sequences as Gaussian parameters μ and σ, which are combined with an auxiliary noise variable ϵ as follows:

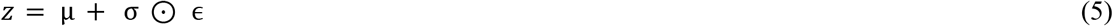

This reparameterized *z* forms the code utilized by the decoder for sequence generation, allowing us to define *p*_θ_ (*z*) as a Gaussian distribution, thereby providing an analytical solution to the gradient of Equation (4)

### Data Representation and Decoding

In our specific implementation, data is represented as a one-hot encoded vector. For a protein of length *L*, an array of shape 23 × *L* is created, where each row contains a 1 in a position corresponding to an amino acid identity, with the remaining row positions containing 0. A total of 23 rows are employed to encode the 20 canonical amino acids, a gap character, and additional less common amino acids. The latent variables *z* are decoded into a Softmax probability distribution with the same array dimensions as the input. The output layer ψ: *R*^23*\*L*^, with each column corresponding to 23 sequence symbols *a ∈ A*, is defined as:

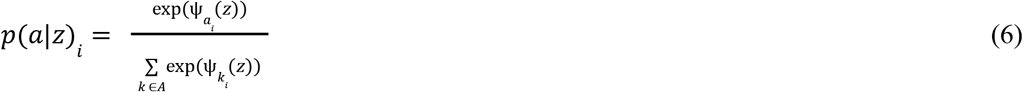

This yields *L* rows with probability values summing to one in each row. The reconstruction error term in Equation (4) evaluates to zero if the input and output matrices are identical, indicating that the only possible sequence at some point *z* is the input sequence.

### Hyperparameters and Training

The model architecture comprises two layers, each of which is an encoder and decoder. For the encoder, the first layer had 2L hidden units while the second layer had L hidden units, where L is the sequence length. Conversely, the decoder’s first layer contained L hidden units, and the second layer had 2L hidden units. ReLU activation functions were employed throughout. The Adam optimizer was utilized with a learning rate of 1e-4, and L2 regularization with a penalty of 1e-4 was applied to the hidden units. The training was terminated if the reconstruction loss did not improve for 50 consecutive epochs. This model, featuring two latent encoding dimensions, was trained on workstations equipped with NVIDIA A100 GPUs.

### Landscape generation

For scoring the generated sequences by VAE, we used the Direct Coupling Analysis(DCA) method, which models protein sequences S of length L via Boltzmann distribution.

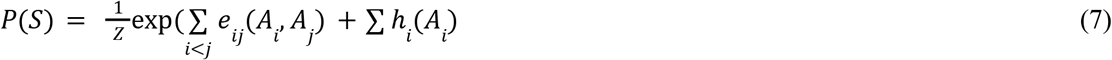

The inferred parameters *e* _*ij*_ encode pairwise coupling between MSA positions and *h*_*i*_ the encoding frequency of amino acids at that position. In a grid-like fashion, equally spaced coordinates of latent space are fed into the VAE decoder to generate a decoded Softmax distribution. The maximum probability sequence from this output is generated as the final sequence. The generated sequence is then given a Hamiltonian score using the parameters obtained from Boltzmann-like DCA distribution, defined as follows:

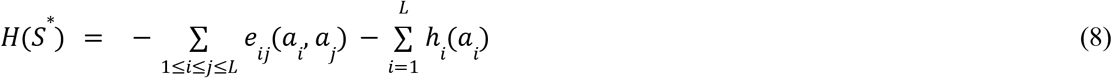

where *S*^*^ = *a*_*i*_ ……. *L* and 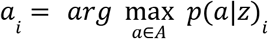

### Protein Structure Prediction with AlphaFold

The initial step in our methodology involves predicting the wild-type (WT) and mutant protein structures based on their sequences using ColabFold, an open-source alternative to AlphaFold2. ColabFold leverages the ‘Evoformer’ module, which integrates multiple sequence alignments and equivariant attention architecture to accurately predict the 3D coordinates of all heavy atoms in a given protein sequence. This tool also provides the predicted local-distance difference test (pLDDT) score, which evaluates the prediction’s accuracy on a scale from 0 to 100, with higher scores indicating better performance [37,38](Jumper et al., 2021 Mirdita et al., 2022). Using the default parameters, we imported the previously collected WT and mutant protein sequences into the ColabFold model to obtain the corresponding PDB files. This process facilitated the generation of high-confidence structural models essential for our subsequent analyses.

### Hydrophobic Contacts and Network Analysis

To calculate interaction fingerprints (IFPs) from the molecular dynamics (MD) simulation and structures generated from Alphafold, we utilized ProLIF (v1.1.0) in conjunction with RDKit (v2021.03.5) and MDAnalysis (v2.4.0), following the procedures outlined in the ProLIF documentation.(David et al., 2021 Landrum et al., 2021 Michaud-Agrawal et al., 2011 Gowers et al., 2016).This setup enabled the extraction of detailed interaction patterns between molecular structures.

#### Hydrophobic Contact Analysis in Static Structures

The CSV files generated from IFP calculations were further processed to evaluate hydrophobic contacts. For static structures, the total number of hydrophobic contacts was normalized by sequence length, yielding per residue hydrophobic contact metrics. This normalization was crucial for comparing proteins of varying lengths generated by the variational autoencoder (VAE).

#### Hydrophobic Contacts in MD Simulations

For the MD simulations, hydrophobic contacts were analyzed by dividing the total count by the frame length. This division helped distinguish between permanent and transient hydrophobic interactions, providing insights into the stability and behavior of the protein under simulated physiological conditions.

### Molecular Dynamics Simulations

The 3D structures generated by AlphaFold were utilized as the initial models for the molecular dynamics (MD) simulations conducted using OpenMM[39]. The amber14-all force field was applied to model the proteins, while the tip3pfb model was used for the water molecules to create the solvated systems [40,41]. Each solvated system underwent energy minimization and equilibration in the NVT ensemble. This was followed by a productive MD run in the NPT ensemble, conducted for 1.2 microseconds at 298 K. During the simulations, a Langevin Middle integrator was employed to maintain a constant temperature of 298 K, and OpenMM’s Monte Carlo barostat ensured a constant pressure of 1 atm[39,42].

#### Structural Analysis and Convergence Assessment

The root mean square deviation (RMSD) of the backbone atom positions and the root mean square fluctuation (RMSF) of individual residues were calculated using MDAnalysis [43,44]. Convergence of the simulations was assessed by comparing RMSF values at different time steps, ensuring fluctuations remained stable as the simulation progressed from shorter to longer time frames. Additionally, minimal changes in the RMSD of the backbone structure after 200 ns indicated stable structural behavior throughout the simulation duration.

## Results

### Evolutionary landscape of Malate Dehydrogenase and its connection with structural diversity

Analysis of the landscape conducted by[32],[45], and [46], has shown that protein sequences trained with a Variational Autoencoder (VAE) tend to get mapped into the low dimensional manifold where functionally viable variants are grouped in distinct basins. VAE encoding also captures the most significant structural variations. We use this insight to elucidate the driving forces of thermal adaptation of cytosolic malate dehydrogenase (cMDH) by utilizing a combination of VAE and Direct Coupling Analysis (DCA) (Fig 2A).

**Figure 2:**
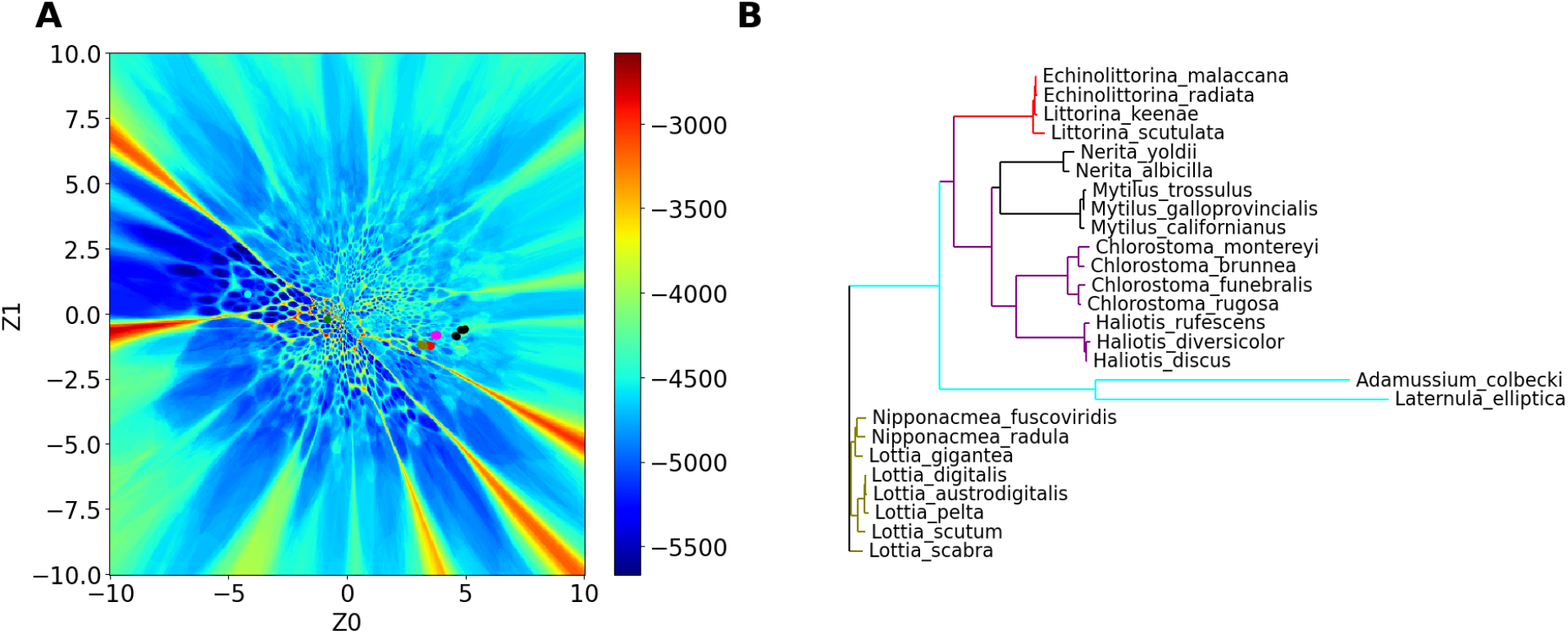
Clustering in the MDH Evolutionary Landscape. (A) This evolutionary landscape plot visualizes the distribution of Direct Coupling Analysis (DCA) scores for cytosolic malate dehydrogenase (cMDH) sequence variants generated by a Variational Autoencoder (VAE). The color gradient, ranging from blue to red, indicates DCA scores, with red representing regions of higher scores. (B) Phylogenetic tree for cMDH, generated using Clustal Omega, is constructed from a distance matrix derived through pairwise sequence comparisons.

The evolutionary landscape mapped by a combination of VAE and DCA allows us to visualize the functional diversity within the MDH family. Characterized by distinct regions demarcated by high and low DCA Hamiltonian values, the landscape reveals the contributions from co-evolved residues. Regions with lower Hamiltonian values are predominantly populated by experimental structures, which correlate with enhanced functional stability due to highly co-evolved residues[47–51]. Within the evolutionary landscape (Fig 2A), species such as *Adamussium colbecki*, with a lethal temperature of 4°C, and Laternula elliptica, with a lethal temperature of 8.3°C, are located in distinct low Hamiltonian basins. This spatial arrangement starkly contrasts with sequences from species enduring higher lethal temperatures, ranging from 32°C to 56°C, which cluster tightly on the opposite side. To assess the pairwise sequence distances among 26 cMDH homologs, we constructed a Guide tree(Fig 2B) using Clustal Omega. This tree, based on a distance matrix derived from pairwise sequence comparisons, highlights distinct clustering patterns. Specifically, species that endure higher lethal temperatures exhibit greater sequence similarity, forming a tighter cluster within the tree. Such clustering highlights greater sequence similarity among high-temperature cMDH orthologs as compared to their low-temperature counterparts, reinforcing the link between sequence homology and adaptability to different thermal conditions[52].

To explore structural diversity in the evolutionary landscape, we analyzed 400 sequences, selecting 100 random sequences covering all major energy basins on the landscape (Figure 3A). Structures were predicted using AlphaFold, and their structural similarity was quantified through Root Mean Square Deviation (RMSD). We used a threshold of 2Å RMSD to classify structures as similar to the known orthologs[53–55]. Four types of structures, clustered according to their RMSD, were identified in the landscape. Homology analysis, by using Interpro[56], revealed two clusters corresponding to cMDH; one cluster included sequences with an active site, similarly positioned as in experimental cMDH structures, while the other lacked this active site.

**Figure 3:**
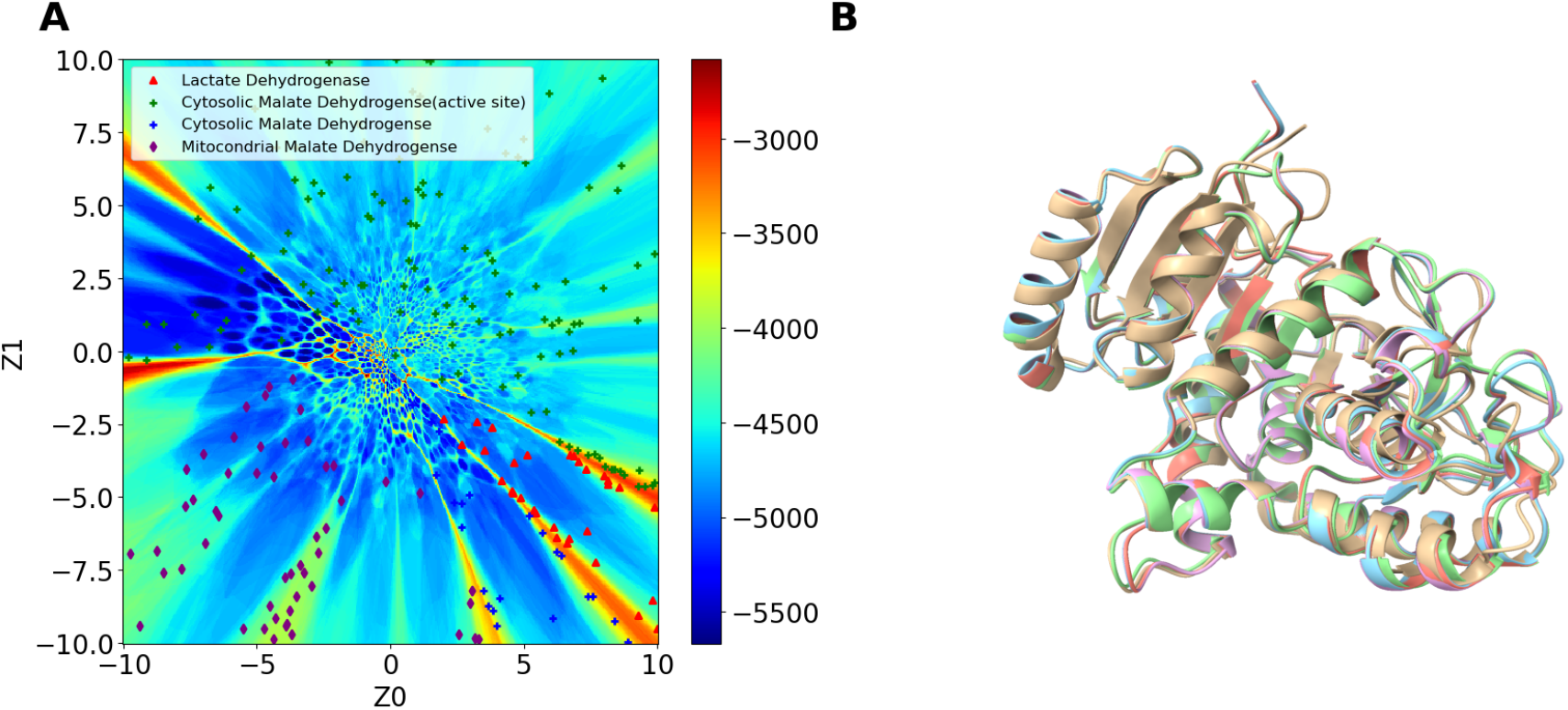
Generated Structures from the Evolutionary Landscape. Scatter points represent the positions of generated structures within the evolutionary landscape. (A) Scatter points indicating the positions of generated structures in the landscape, where four types of structures (cMDH with active site, cMDH without active site, lactate dehydrogenase (LDH), and mitochondrial MDH) are clustered based on their RMSD values. (B) Cytosolic malate dehydrogenase (cMDH) structures aligned with generated sequence structures, all with root mean square deviation (RMSD) < 2.0 Å.

Additionally, one cluster was identified as mitochondrial malate dehydrogenase and another as lactate dehydrogenase. Intriguingly, the landscape also reveals several structurally similar sequences near the borders of high DCA energy barriers, indicating a distinct boundary in structural adaptation. Moreover, no sequences with similar structures were found across these barrier regions, underscoring a significant demarcation in the evolutionary landscape that correlates with structural and functional divergence from cMDH.

To gain microscopic insight into the evolution of the thermal stability of *Malate Hydrogenase* we calculated hydrophobic contacts using the structures generated from the VAE landscape (Fig 4). Hydrophobic contacts were calculated per residue based on atomistic distances grouped into residues. Due to the variability in sequence length, which includes gaps of insertion and deletion similar to those found in multiple sequence alignment files, the total number of hydrophobic contacts was normalized by dividing by the total number of residues. This normalization process allows for consistently comparing hydrophobic contacts per residue across the landscape. We uncovered an inverse correlation between Hamiltonian values and hydrophobic contact per residue (Fig 4A). Since hydrophobic contacts are integral to the structural stability of proteins, particularly at higher temperatures, this trend suggests that structural stability diminishes as sequences shift towards higher DCA Hamiltonian values[57,58].

**Figure 4:**
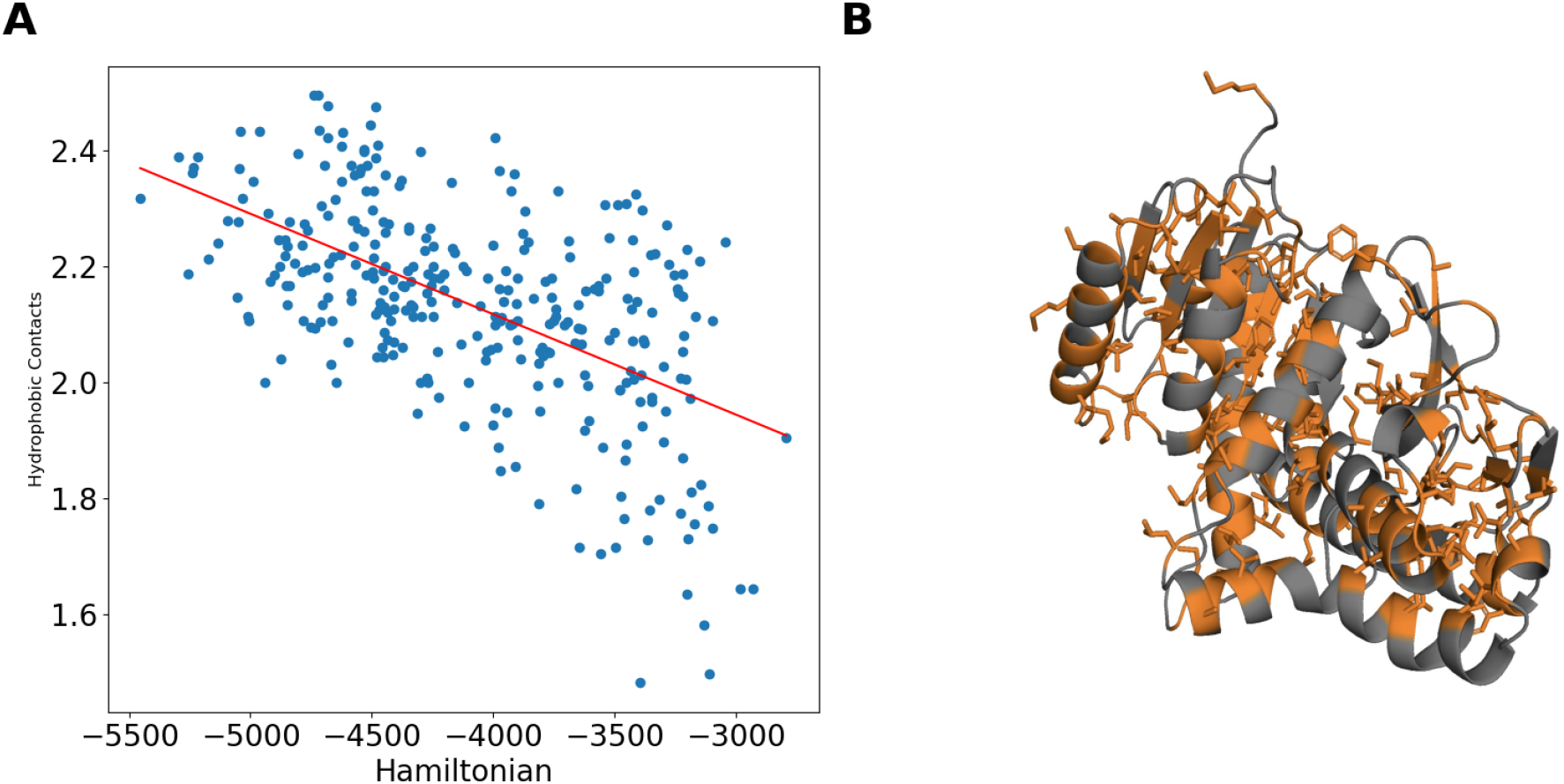
Relationship Between DCA Hamiltonian and Normalized Hydrophobic Contacts in Protein Structures. (A) This scatter plot depicts the correlation between DCA Hamiltonian values and the normalized number of hydrophobic contacts per residue across 400 protein structures generated from the VAE landscape. The red trend line indicates a decreasing number of hydrophobic contacts as the DCA Hamiltonian, plotted on the x-axis, increases. (B) Hydrophobic residue in the generated structure.

### Dynamics sheds light on evolving thermal activity in cMDH homologs

While structures provide clues on thermal adaptation, it is, without doubt, the temperature-dependent dynamics of enzymes that report most accurately on catalytic activities[59,60]. To this end, we have conducted microsecond-long atomistic molecular dynamics simulations at room temperature and quantified enzyme dynamics via mean square fluctuation (RMSF) of contacts and cartesian displacements. We identify two regions (Figure 6B) with significant fluctuations: residues 91–105, forming the catalytic loop folding onto the catalytic site during ligand binding (highlighted in red in Figure 6A), and residues 230–245, which are involved in the catalytic process (highlighted in blue in Figure 6A)[7]. Notably, *Adamussium Colbecki*, adapted to colder environments, exhibits the highest RMSF, consistent with its low lethal temperature of approximately 277K. While fluctuations in the first mobile region remained consistent across species, notable differences were observed in the dynamics of the second region as species adaptation ranged from colder to warmer environments(Figure 6B). This observation indicates that although the dynamic behavior of the catalytic area remains consistent, the distinct peaks observed in Region 2, which is also involved in catalysis, should be subject to further study. Hydrophobic interactions, salt bridges, van der Waals forces, and aromatic stacking (cite related papers) are key contributors to thermal stability. However, the precise mechanism by which these interactions collectively influence thermal stability remains poorly understood (Supplementary Fig.1). A protein must maintain sufficient stability to preserve its structure while remaining flexible enough to carry out its biological functions near its optimal operating temperature [61–63]. The importance of balancing stability and degree of functional dynamism [64–66] has led us to investigate both permanent and transient hydrophobic interactions throughout the simulation. Permanent hydrophobic contacts, defined as those present for more than 95% of the simulation time, were contrasted with transient contacts, which occur for less than 50%. Interestingly, the frequency of transient hydrophobic contacts decreases in enzymes adapted to higher temperatures (Fig S2).

**Figure 6:**
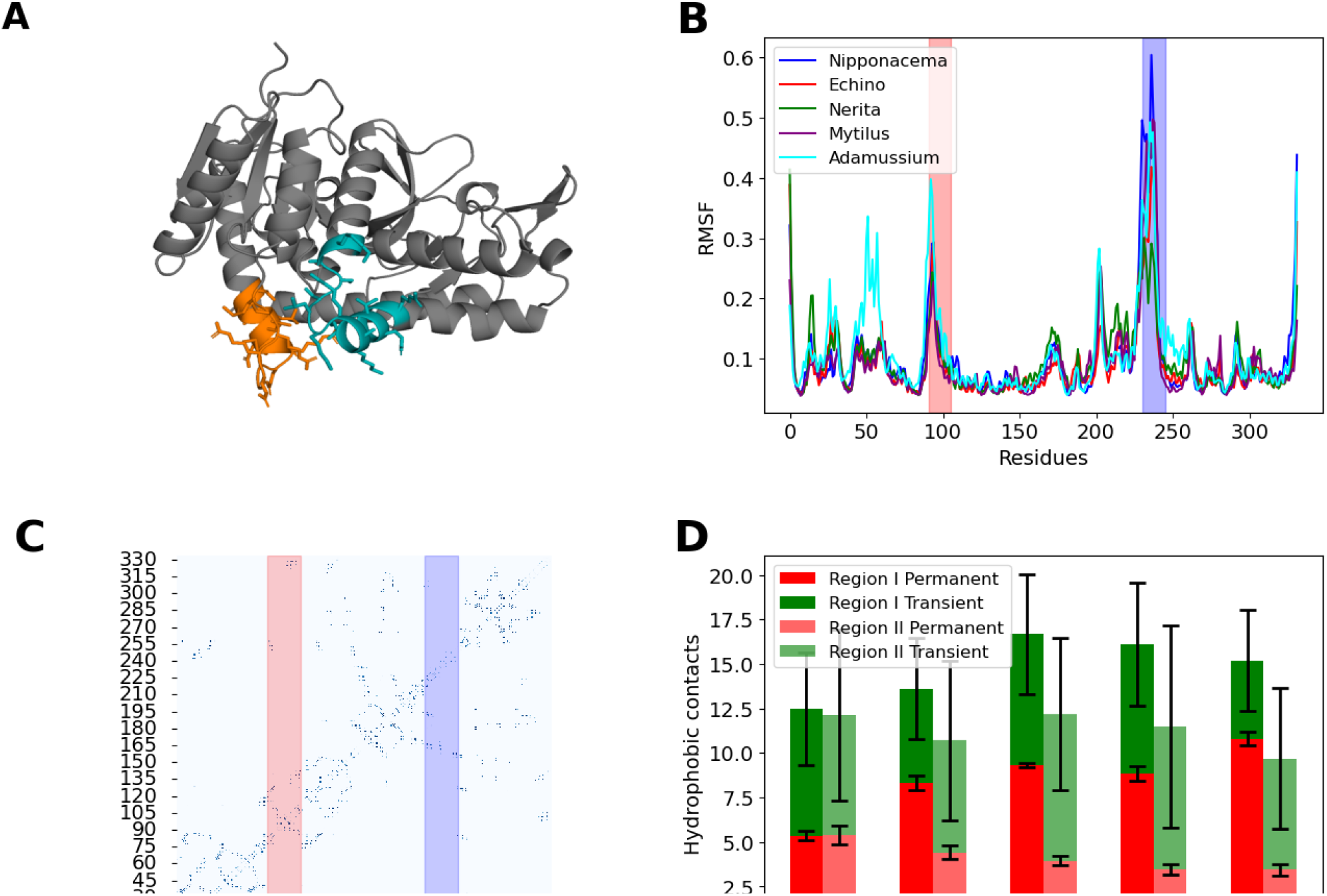
Molecular Dynamics Analysis of Side Chain Movements (RMSF) of cMDH Across Various Species at ambient temperature. (A) Structure of cytosolic malate dehydrogenase (cMDH) highlighting functionally active residues. The catalytic loop (residues 91–105) is shown in red, while the additional catalytic region (residues 230–245) is depicted in blue. (B) Root Mean Square Fluctuation (RMSF) profiles at 298K for cMDH from five species: *Echinolittorina malaccana, Nerita yoldi, Mytilus californianus, Adamussium colbecki*, and *Nipponacmea radula*. Peaks in RMSF at residues 91–105 and 230–245, marked in red and blue, respectively, indicate regions of significant fluctuation. (C) The contact map highlights that the region containing catalytically active residues forms long-range hydrophobic contacts. (D) Permanent and transient hydrophobic contacts in Region I (residues 80–115) and Region II (residues 225–250) across different cMDH families, with psychrophilic enzymes on the left and thermophilic enzymes on the right, emphasizing differences in contact dynamics related to thermal adaptation.

We found that the catalytically significant residues (90-105 and 225-245) showed an absence of permanent hydrophobic contacts. This prompted us to explore the dynamic hydrophobic interactions near the catalytic and highly flexible domain boundaries, focusing on residues 80-110 and 220-250. Notably, in addition to being catalytically active, these regions in cMDH are responsible for long-range hydrophobic interactions (Figure 6C).

We find that cMDH homologs from warmer climates exhibit a marked increase in permanent hydrophobic contacts (Figure 6D) within Region I (residues 80-115) compared to those from colder environments. On the other hand, we find a decrease in the frequency of permanent hydrophobic contacts in Region II (residues 225-250) at the same temperature ranges (Figure 7B). Interestingly, Region I, a key catalytic area, is predominantly characterized by permanent hydrophobic contacts (Figure 6D). In contrast, Region II, also critical for catalysis, primarily displays transient hydrophobic contacts corresponding to regions of high fluctuation. Moreover, only *Adamussium*, an enzyme with a lethal temperature of 4°C, shows more transient frequency than permanent contacts in Region I.. In contrast, mesophilic and thermophilic enzymes show a higher frequency of preserved hydrophobic contacts in this region during the simulation, suggesting the role of preserved interactions in the thermal adaptation of these enzymes.[67].

**Figure 7:**
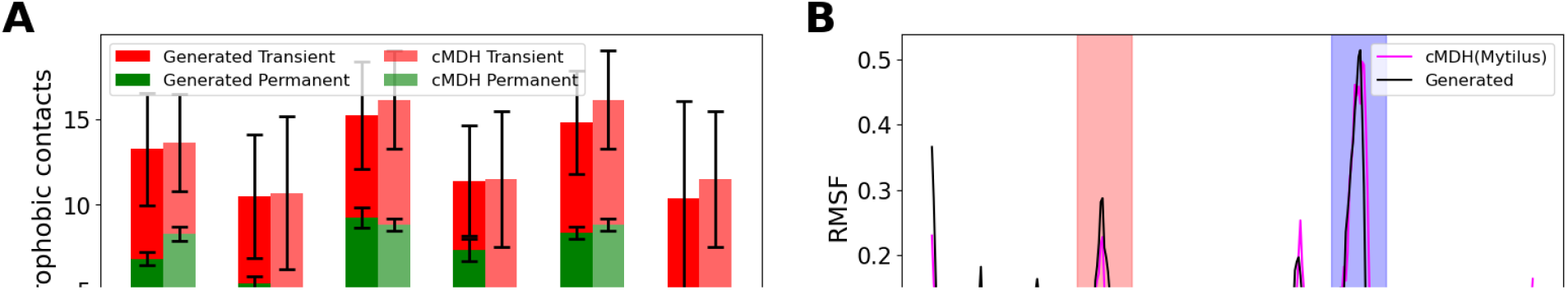
Dynamic hydrophobic contact Molecular Dynamic analysis of side chain movements (RMSF) for Generated and Known cMDH Structures: (A) Figure shows the comparison between transient(red) and permanent(green) hydrophobic contacts (B,C,D) Root Mean Square Fluctuations (RMSF) of computationally generated cMDH structures compared to known cMDH structures from various species, including *Mytilus californianus, Nerita yoldii*, and *Echinolittorina radita*. Each graph shows the RMSF for a generated structure (blue) alongside the RMSF for a known cMDH structure(red), clustered at that location, highlighting the close correspondence in dynamic properties.

These findings highlight the crucial role of dynamic hydrophobic interactions within specific regions, potentially enhancing the enzyme’s thermal adaptability by modulating the flexibility of areas critical to catalysis. The observed increase in permanent hydrophobic contacts in particular regions suggests a mechanism through which these cMDH stabilize their structure under elevated temperatures, thereby compensating for increased thermal motion.

The RMSF profiles (Figs. 7B-D) of the generated structures, derived from the cMDH clustering locations in the sequence landscape, closely resemble those of known cMDH structures from species such as *Mytilus californianus* and *Nerita yoldii*. We examined the dynamic hydrophobic contact frequency to explore the reasons for similarities and differences in the flexibility of catalytic regions. We observed that Region I exhibits similar flexibility between the generated and cMDH structures, as reflected in the RMSF plots (Figs. 7B-D). This can be correlated with Figure 7B, where Region I shows a similar frequency of both permanent and transient hydrophobic contacts, indicating structural conservation in this catalytic region.

In contrast, Region II shows differences in flexibility, which can also be explained by dynamic contact frequency analysis. For instance, comparing the dynamic hydrophobic contact frequency of *Nerita yoldii* with a generated structure reveals greater flexibility in Region II for the latter (Figure 7C), and lower flexibility for *Nerita yoldii* (Figure 7D). Our analysis shows that structures with lower fluctuations have a higher frequency of permanent hydrophobic contacts and fewer transient ones, while the more flexible generated structure displays the opposite trend. Dynamic contact analysis in this case revealed that higher permanent hydrophobic contacts correlate with reduced flexibility, further confirming this relationship.

We also examined cases where fluctuations were comparable (Fig 7B) such as the case in *Mytilus californianus* which shows a dynamic contact frequency similar to the generated structure. This parallel in dynamic properties between the known and generated structures supports the validity of our computational models as accurate representations of the real dynamical behavior of cMDH enzymes [68]. Additionally, this analysis highlights the importance of dynamic hydrophobic contact frequency in understanding protein flexibility, with more permanent contacts correlating with less flexibility and transient contacts correlating with greater flexibility.

### Dynamics of sequence evolution using generated structures from evolutionary landscape

To investigate evolutionary patterns of conformational dynamics across different cMDH clusters of homologs, we conducted a detailed study along a path from the *Echinolittorina* family, clustered in the evolutionary landscape, to the *Mytilus* family, with the *Nerita* family cluster present along this path (Fig 9B). We selected eight representative structures from this trajectory. We subjected them to all-atom simulations over a microsecond at room temperature, maintaining the same conditions as those used for the cMDH enzymes. Consistent with our general observations for cMDH enzymes, the RMSF profiles for sampled structures (Figure 9) showed similar fluctuations across most regions, except for Region 2, where each structure exhibited unique peaks. The fluctuations near the catalytic regions remained constant for all homologs, highlighting the structural conservation in these functionally important regions.

**Figure 9:**
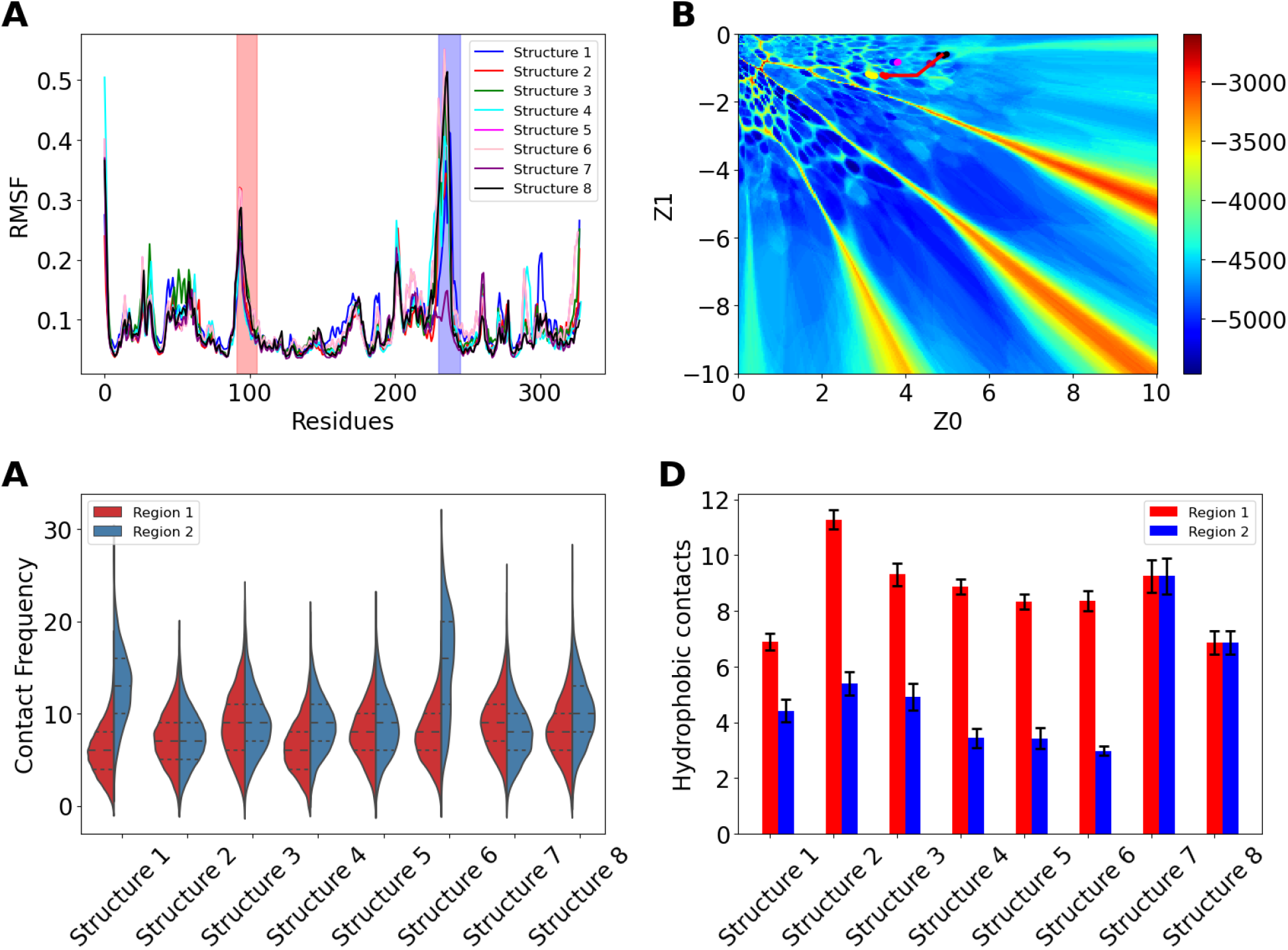
Change in the dynamics and thermal stability along a path connecting cMDH functional clusters. (A) Root Mean Square Fluctuation (RMSF) profiles at 298K for eight generated structures along a trajectory connecting the *Echinolittorina, Nerita, and Mytilus* family. (B) The path between families in the evolutionary landscape. (C) Plot showing transient hydrophobic contacts in both regions, residue 80-110 and residue 225-250 (D) Plot showing permanent hydrophobic contacts in both regions, residue 80-110 and residue 225-250

One of the eight generated structures (Figure 9D) exhibits the highest RMSF in Region II and has the highest number of transient hydrophobic contacts and the lowest number of permanent hydrophobic contacts. Conversely, a different structure, depicted as structure 7 (Figure 9D) with the lowest RMSF, shows the highest number of permanent hydrophobic contacts and one of the lowest counts of transient hydrophobic contacts in Region II. All generated structures demonstrate more permanent hydrophobic contacts in Region I (Figure 9C) than transient contacts, aligning with the cMDH hydrophobic contact patterns (Figure 7A). These findings underscore the importance of maintaining a higher count of permanent hydrophobic contacts relative to transient ones to preserve catalytic activity at elevated temperatures.

## Discussion

Despite decades of research on cytosolic malate dehydrogenase (cMDH), detailed insights into how these enzymes adapt to varying thermal environments to catalyze the conversion of malate to oxaloacetate have yet to be discovered. In this study, we employed a combination of AI, bioinformatics, and molecular simulation techniques to generate an evolutionary landscape for cMDH, providing a novel framework for understanding thermal adaptation at the molecular level. Specifically, we focused on dynamic contact variability, a critical factor in maintaining enzyme stability across different temperatures.

Using multiple sequence alignments (MSA) and a variational autoencoder (VAE), we generated a diverse sequence landscape comprising approximately 250,000 potential cMDH sequences. Selection of these sequences based on Direct Coupling Analysis (DCA) scores, Hamiltonian barriers, and clustering patterns allowed us to infer critical structural and functional properties. Notably, sequences clustered with low Hamiltonian scores demonstrated a strong correlation with enhanced structural stability, offering deeper insights into the relationship between sequence space and protein function.

Our structural analyses revealed a significant inverse relationship between Hamiltonian scores and the number of hydrophobic contacts per residue. Lower Hamiltonian scores were associated with reduced hydrophobic contacts, which, as supported by prior studies, correlates with decreased thermal stability. This finding underscores the importance of hydrophobic interactions in the thermal adaptation of cMDH, with dynamic contact variability serving as a key mechanism for stabilizing enzyme structures under different thermal conditions.

The dynamic analysis of the generated sequences was elucidated via microsecond-long atomistic simulations, providing further insight into the evolutionary landscape of cMDH. The dynamic behavior of these sequences closely mirrored that of experimentally characterized cMDH structures from marine mollusks. Consistent with previous research, we confirmed that cMDH possesses two mobile regions essential for catalysis. Our results demonstrate that the variability in dynamics across species adapted to different temperature ranges is largely driven by changes in the number of dynamic hydrophobic interactions near these mobile regions, influenced by specific mutations as one moves from one species to another.

In conclusion, our study highlights the pivotal role of hydrophobic interactions, sequence variability, and dynamic flexibility in the thermal adaptation of cMDH enzymes. By leveraging a deep learning-generated sequence landscape and integrating structural and dynamic analyses, we provide new insights into the relationship between sequence, structure, and thermal stability. Our findings clarify how mutations drive adaptation across species, enhancing our understanding of cMDH stability in diverse thermal environments. This work underscores the potential of AI-driven approaches in uncovering the molecular mechanisms of thermal adaptation and opens new avenues for exploring similar adaptations in other protein systems. These insights have significant implications for the rational design of enzymes with tailored thermal properties, contributing to biotechnology and protein engineering.

## References

1. Nava G, Laclette JP, Bobes R, Carrero JC, Reyes-Vivas H, Enriquez-Flores S, et al. Cloning, sequencing and functional expression of cytosolic malate dehydrogenase from Taenia solium: Purification and characterization of the recombinant enzyme. Exp Parasitol. 2011;128: 217–224.

2. Minárik P, Tomásková N, Kollárová M, Antalík M. Malate dehydrogenases--structure and function. Gen Physiol Biophys. 2002;21: 257–265.

3. Goto Y, Fink AL. Conformational states of beta-lactamase: molten-globule states at acidic and alkaline pH with high salt. Biochemistry. 1989;28: 945–952.

4. Yueh AY, Chung CS, Lai YK. Purification and molecular properties of malate dehydrogenase from the marine diatom Nitzschia alba. Biochem J. 1989;258: 221–228.

5. Chao Y-C, Merritt M, Schaefferkoetter D, Evans TG. High-throughput quantification of protein structural change reveals potential mechanisms of temperature adaptation in Mytilus mussels. BMC Evol Biol. 2020;20: 28.

6. Pucci F, Rooman M. Physical and molecular bases of protein thermal stability and cold adaptation. Curr Opin Struct Biol. 2017;42: 117–128.

7. Dong Y-W, Liao M-L, Meng X-L, Somero GN. Structural flexibility and protein adaptation to temperature: Molecular dynamics analysis of malate dehydrogenases of marine molluscs. Proc Natl Acad Sci U S A. 2018;115: 1274–1279.

8. Dong M. A Minireview on Temperature Dependent Protein Conformational Sampling. Protein J. 2021;40: 545–553.

9. Zhang J, Liu S, Chen M, Chu H, Wang M, Wang Z, et al. Unsupervisedly prompting AlphaFold2 for accurate few-shot protein structure prediction. J Chem Theory Comput. 2023;19: 8460–8471.

10. Siddiqui KS, Cavicchioli R. Cold-adapted enzymes. Annu Rev Biochem. 2006;75: 403–433.

11. Szilágyi A, Závodszky P. Structural differences between mesophilic, moderately thermophilic and extremely thermophilic protein subunits: results of a comprehensive survey. Structure. 2000;8: 493–504.

12. Vieille C, Zeikus GJ. Hyperthermophilic enzymes: sources, uses, and molecular mechanisms for thermostability. Microbiol Mol Biol Rev. 2001;65: 1–43.

13. Somero GN. Proteins and temperature. Annu Rev Physiol. 1995;57: 43–68.

14. Feller G, Gerday C. Psychrophilic enzymes: hot topics in cold adaptation. Nat Rev Microbiol. 2003;1: 200–208.

15. DePristo MA, Weinreich DM, Hartl DL. Missense meanderings in sequence space: a biophysical view of protein evolution. Nat Rev Genet. 2005;6: 678–687.

16. Tokuriki N, Tawfik DS. Protein dynamism and evolvability. Science. 2009;324: 203–207.

17. Bloom JD, Arnold FH. In the light of directed evolution: Pathways of adaptive protein evolution. Proceedings of the National Academy of Sciences. 2009;106: 9995–10000.

18. Morcos F, Pagnani A, Lunt B, Bertolino A, Marks DS, Sander C, et al. Direct-coupling analysis of residue coevolution captures native contacts across many protein families. Proc Natl Acad Sci U S A. 2011;108: E1293–301.

19. Marks DS, Colwell LJ, Sheridan R, Hopf TA, Pagnani A, Zecchina R, et al. Protein 3D structure computed from evolutionary sequence variation. PLoS One. 2011;6: e28766.

20. Hopf TA, Schärfe CPI, Rodrigues JPGLM, Green AG, Kohlbacher O, Sander C, et al. Sequence co-evolution gives 3D contacts and structures of protein complexes. Elife. 2014;3. doi:10.7554/eLife.03430

21. Ovchinnikov S, Kamisetty H, Baker D. Robust and accurate prediction of residue–residue interactions across protein interfaces using evolutionary information. Elife. 2014;3: e02030.

22. Michel M, Hayat S, Skwark MJ, Sander C, Marks DS, Elofsson A. PconsFold: improved contact predictions improve protein models. Bioinformatics. 2014;30: i482–8.

23. Morcos F, Hwa T, Onuchic JN, Weigt M. Direct coupling analysis for protein contact prediction. Methods Mol Biol. 2014;1137: 55–70.

24. Weigt M, White RA, Szurmant H, Hoch JA, Hwa T. Identification of direct residue contacts in protein–protein interaction by message passing. Proceedings of the National Academy of Sciences. 2009;106: 67–72.

25. Ekeberg M, Hartonen T, Aurell E. Fast pseudolikelihood maximization for direct-coupling analysis of protein structure from many homologous amino-acid sequences. J Comput Phys. 2014;276: 341–356.

26. Brookes DH, Park H, Listgarten J. Conditioning by adaptive sampling for robust design. arXiv [cs.LG]. 2019. Available: http://arxiv.org/abs/1901.10060

27. Linder J, Bogard N, Rosenberg AB, Seelig G. A generative neural network for maximizing fitness and diversity of synthetic DNA and protein sequences. Cell Syst. 2020;11: 49–62.e16.

28. Du X, Sun S, Hu C, Yao Y, Yan Y, Zhang Y. DeepPPI: Boosting Prediction of Protein–Protein Interactions with Deep Neural Networks. J Chem Inf Model. 2017;57: 1499–1510.

29. Zou X, Wang G, Yu G. Protein Function Prediction Using Deep Restricted Boltzmann Machines. Biomed Res Int. 2017;2017: 1729301.

30. Ingraham J, Garg VK, Barzilay R, Jaakkola T. Generative models for graph-based protein design. 2019; 15794–15805.

31. Riesselman AJ, Ingraham JB, Marks DS. Deep generative models of genetic variation capture the effects of mutations. Nat Methods. 2018;15: 816–822.

32. Ziegler C, Martin J, Sinner C, Morcos F. Latent generative landscapes as maps of functional diversity in protein sequence space. Nat Commun. 2023;14: 2222.

33. Eddy SR. Accelerated Profile HMM Searches. PLoS Comput Biol. 2011;7: e1002195.

34. Finn RD, Clements J, Eddy SR. HMMER web server: interactive sequence similarity searching. Nucleic Acids Res. 2011;39: W29–37.

35. Sievers F, Wilm A, Dineen D, Gibson TJ, Karplus K, Li W, et al. Fast, scalable generation of high-quality protein multiple sequence alignments using Clustal Omega. Mol Syst Biol. 2011;7: 539.

36. Kingma DP, Welling M. Auto-Encoding Variational Bayes. arXiv [stat.ML]. 2013. Available: http://arxiv.org/abs/1312.6114v11

37. Jumper J, Evans R, Pritzel A, Green T, Figurnov M, Ronneberger O, et al. Highly accurate protein structure prediction with AlphaFold. Nature. 2021;596: 583–589.

38. Mirdita M, Schütze K, Moriwaki Y, Heo L, Ovchinnikov S, Steinegger M. ColabFold: making protein folding accessible to all. Nat Methods. 2022;19: 679–682.

39. Eastman P, Swails J, Chodera JD, McGibbon RT, Zhao Y, Beauchamp KA, et al. OpenMM 7: Rapid development of high performance algorithms for molecular dynamics. PLoS Comput Biol. 2017;13: e1005659.

40. Jorgensen WL, Chandrasekhar J, Madura JD, Impey RW, Klein ML. Comparison of simple potential functions for simulating liquid water. J Chem Phys. 1983;79: 926–935.

41. Wang L-P, Martinez TJ, Pande VS. Building force fields: An automatic, systematic, and reproducible approach. J Phys Chem Lett. 2014;5: 1885–1891.

42. Martyna GJ, Tobias DJ, Klein ML. Constant pressure molecular dynamics algorithms. J Chem Phys. 1994;101: 4177–4189.

43. Michaud-Agrawal N, Denning EJ, Woolf TB, Beckstein O. MDAnalysis: a toolkit for the analysis of molecular dynamics simulations. J Comput Chem. 2011;32: 2319–2327.

44. Gowers R, Linke M, Barnoud J, Reddy T, Melo M, Seyler S, et al. MDAnalysis: A python package for the rapid analysis of molecular dynamics simulations. Proceedings of the Python in Science Conference. SciPy; 2016. pp. 98–105.

45. Tian H, Jiang X, Trozzi F, Xiao S, Larson EC, Tao P. Explore Protein Conformational Space With Variational Autoencoder. Front Mol Biosci. 2021;8: 781635.

46. Detlefsen NS, Hauberg S, Boomsma W. Learning meaningful representations of protein sequences. Nat Commun. 2022;13: 1914.

47. Dimas RP, Jiang X-L, Alberto de la Paz J, Morcos F, Chan CTY. Engineering repressors with coevolutionary cues facilitates toggle switches with a master reset. Nucleic Acids Res. 2019;47: 5449–5463.

48. Russ WP, Figliuzzi M, Stocker C, Barrat-Charlaix P, Socolich M, Kast P, et al. An evolution-based model for designing chorismate mutase enzymes. Science. 2020;369: 440–445.

49. Cheng RR, Nordesjö O, Hayes RL, Levine H, Flores SC, Onuchic JN, et al. Connecting the Sequence-Space of Bacterial Signaling Proteins to Phenotypes Using Coevolutionary Landscapes. Mol Biol Evol. 2016;33: 3054–3064.

50. Figliuzzi M, Jacquier H, Schug A, Tenaillon O, Weigt M. Coevolutionary Landscape Inference and the Context-Dependence of Mutations in Beta-Lactamase TEM-1. Mol Biol Evol. 2016;33: 268–280.

51. Bisardi M, Rodriguez-Rivas J, Zamponi F, Weigt M. Modeling Sequence-Space Exploration and Emergence of Epistatic Signals in Protein Evolution. Mol Biol Evol. 2022;39. doi:10.1093/molbev/msab321

52. Liao M-L, Somero GN, Dong Y-W. Comparing mutagenesis and simulations as tools for identifying functionally important sequence changes for protein thermal adaptation. Proc Natl Acad Sci U S A. 2019;116: 679–688.

53. Xin F, Radivojac P. Post-translational modifications induce significant yet not extreme changes to protein structure. Bioinformatics. 2012;28: 2905–2913.

54. Krissinel E. On the relationship between sequence and structure similarities in proteomics. Bioinformatics. 2007;23: 717–723.

55. Lazar T, Guharoy M, Vranken W, Rauscher S, Wodak SJ, Tompa P. Distance-Based Metrics for Comparing Conformational Ensembles of Intrinsically Disordered Proteins. Biophys J. 2020;118: 2952–2965.

56. Paysan-Lafosse T, Blum M, Chuguransky S, Grego T, Pinto BL, Salazar GA, et al. InterPro in 2022. Nucleic Acids Res. 2023;51: D418–D427.

57. Baldwin RL. Temperature dependence of the hydrophobic interaction in protein folding. Proc Natl Acad Sci U S A. 1986;83: 8069–8072.

58. Widom B, Bhimalapuram P, Koga K. The hydrophobic effect. Phys Chem Chem Phys. 2003;5: 3085–3093.

59. Thomas TM, Scopes RK. The effects of temperature on the kinetics and stability of mesophilic and thermophilic 3-phosphoglycerate kinases. Biochem J. 1998;330 (Pt 3): 1087–1095.

60. Daniel RM, Danson MJ. A new understanding of how temperature affects the catalytic activity of enzymes. Trends Biochem Sci. 2010;35: 584–591.

61. Kim SY, Hwang KY, Kim SH, Sung HC, Han YS, Cho Y. Structural basis for cold adaptation. Sequence, biochemical properties, and crystal structure of malate dehydrogenase from a psychrophile Aquaspirillium arcticum. J Biol Chem. 1999;274: 11761–11767.

62. Fitter J, Herrmann R, Dencher NA, Blume A, Hauss T. Activity and stability of a thermostable alpha-amylase compared to its mesophilic homologue: mechanisms of thermal adaptation. Biochemistry. 2001;40: 10723–10731.

63. Folch B, Dehouck Y, Rooman M. Thermo- and mesostabilizing protein interactions identified by temperature-dependent statistical potentials. Biophys J. 2010;98: 667–677.

64. Sanchez-Ruiz JM. Protein kinetic stability. Biophys Chem. 2010;148: 1–15.

65. Karshikoff A, Nilsson L, Ladenstein R. Rigidity versus flexibility: the dilemma of understanding protein thermal stability. FEBS J. 2015;282: 3899–3917.

66. Burns D, Venditti V, Potoyan DA. Temperature-sensitive contact modes allosterically gate TRPV3. bioRxivorg. 2023. doi:10.1101/2023.01.02.522497

67. Noivirt-Brik O, Horovitz A, Unger R. Trade-off between positive and negative design of protein stability: from lattice models to real proteins. PLoS Comput Biol. 2009;5: e1000592.

68. Somero GN. Temperature Adaptation of Enzymes: Biological Optimization Through Structure-Function Compromises. Annu Rev Ecol Syst. 1978;9: 1–29.

